# A conserved superlocus regulates above- and belowground root initiation

**DOI:** 10.1101/2020.11.11.377937

**Authors:** Moutasem Omary, Naama Gil-Yarom, Chen Yahav, Evyatar Steiner, Idan Efroni

## Abstract

During plant post-embryonic growth new meristems and associated stem cells form in different development contexts in order to respond to environmental cues. While underground lateral roots initiate from designated cells in the main root, an unknown mechanism allows cells to bypass the root/shoot identity trajectory and generate shoot-borne-roots. Using single-cell profiling of tomato (*Solanum lycoperiscum)* stems we isolated a rare transient cell population that serve as progenitors for shoot-borne-root meristems. Analysis of this population identified a transcription factor required for the formation of shoot-borne-roots which we named *SHOOT BORNE ROOTLESS (SBRL)*. Evolutionary analysis revealed that *SBRL* function is deeply conserved in angiosperms and that it arose as part of an ancient duplicated superlocus, only lost in root-less plants, containing both shoot-borne and lateral root initiation regulators. We propose that the ability to activate a common transition state with context-specific regulators allows the remarkable developmental plasticity found in plants.

**One Sentence Summary:** Highly conserved superlocus of LBD genes, acting within an early transition identity, regulates shoot-borne and lateral root formation.

## Main Text

The body of vascular plants is divided into the root and shoot systems. Roots are formed via the activity of root meristems, composed of tissue-specific stem cells, or initials, arranged around slowly dividing quiescent cells and protected by the root cap (*1*). The embryonic root and shoot meristems initiate from distinct cells, following the establishment of apical-basal polarity (*2*). Post-embryogenesis, lateral roots (LR) branch off from pre-patterned cells in the primary root pericycle, a specialized tissue that maintains its pluripotency throughout adulthood (*3, 4*). Despite the early root-shoot lineage separation, most plants generate post-embryonic shoot-borne roots (SBR) to produce multiple root systems (*5*). Reflecting the conceptual separation between root and shoot lineages, these roots are commonly referred to as ‘adventitious’ (“in the wrong place”) (*6*). However SBR arise as part of the normal development of many plants and fossil evidence suggests that root-bearing shoots were the dominant body plan of early angiosperms (*7*). Therefore, classical botanists made a distinction between ‘adventitious’ roots, like wound-induced roots, and naturally-occurring SBR (*5, 8*). Although the ability to naturally form such roots is a hallmark of plant developmental plasticity, little information is available on the development of these organs. Currently, the nature of their relation to the LR and the identity of their tissue of origin remain disputed (*9*).

Two key plant hormones, auxin and cytokinin, are vital for the regulation of root initiation. Auxin response is activated in LR meristem progenitors, while response to cytokinin marks the LR’s flanking cells (*3, 10*). Auxin triggers multiple processes during LR initiation, including promotion of cell growth and mitotic cell division via the activation of *LATERAL ORGAN BOUNDARIES DOMAIN (LBD)* transcription factors (*11–15*). Other auxin-induced transcription factors, such as members of the *PLETHORA/AINTEGUMENTA* family (*16*), are required for the establishment of proper cell division patterns and, at later stages, for the acquisition of specific cell fates (*17, 18*). Scarce genetic and transcriptomic evidence from monocots, where SBR form the bulk of the root system, suggests that similar gene families play a role in both SBR and LR initiation (*9, 19*). Thus, in maize and rice, SBR initiation requires the activity of the auxin-induced *LBD* transcription factors *ROOTLESS CONCENRNING CROWN AND SEMINAL ROOT* (*RTCS*) and *CROWN ROOTLESS1* (*CRL1*) (*20–22*), as well as the *PLETHORA/AINTEGUMENTA* (PLT) family protein *CRL5* (*23*). Seemingly unique to SBR, reduced expression of the WUSCHEL-like protein WOX11 results in a delay in SBR development (*24*). The common model plant *Arabidopsis thaliana* lacks a post-embryonic SBR system. Thus, to study SBR initiation in dicots at high resolution, we turned to tomato (*Solanum lycoperiscum*), a vine that naturally generates a large number of SBR from easily accessible stems (Fig. 1, A to F).

**Fig. 1.**
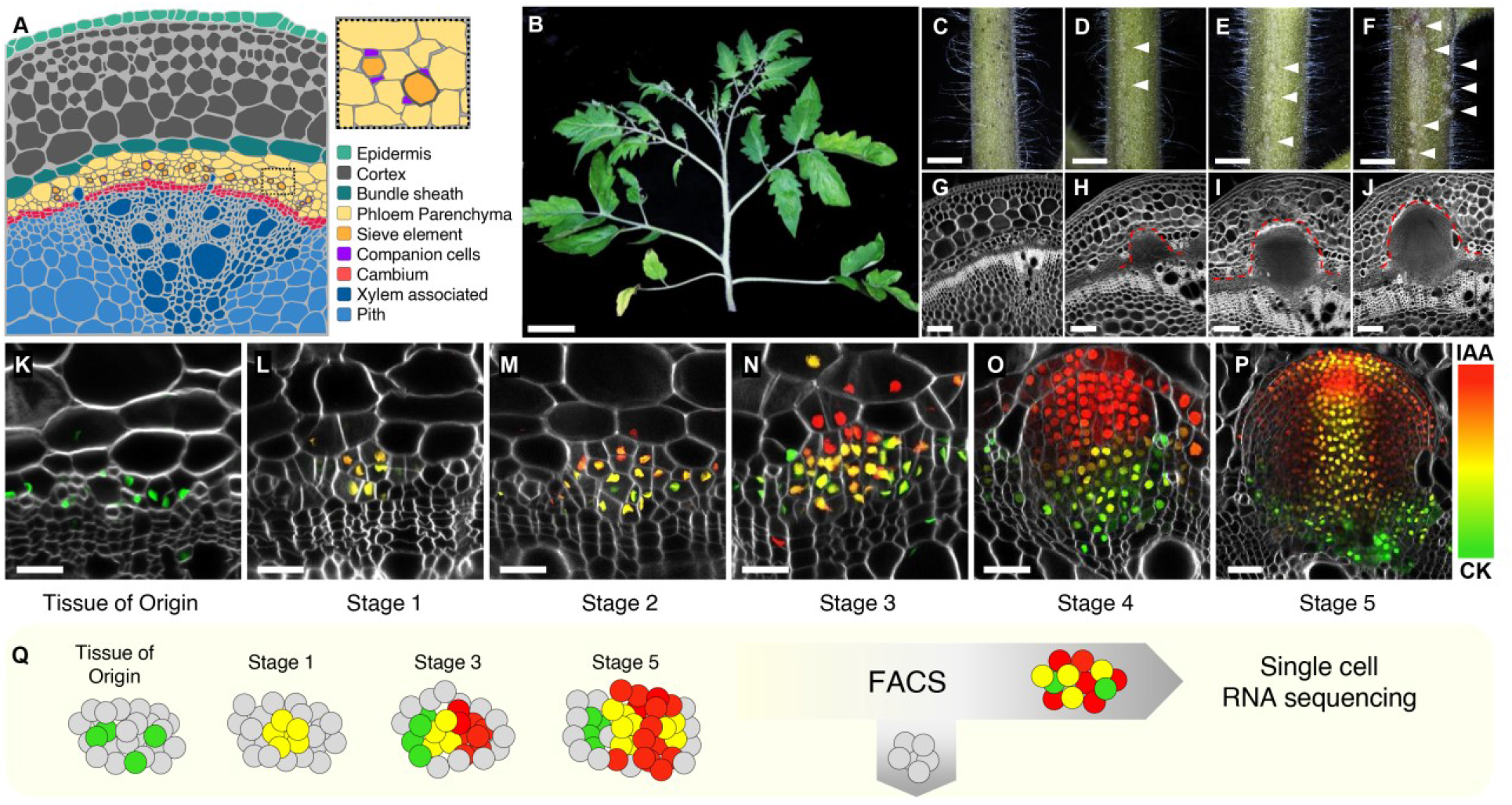
SBR formation in tomato. (**A**) 2D illustration of the different cell types in a tomato stem section and their spatial organization. Inset shows a magnification of the phloem region. (**B**) A shoot of an 8-week-old tomato plant. **(C** to **J)** Close up (**C** to **F**) and confocal images of cross-sections (**G** to **J**) of four internodes of the plant in (B), from young (**C** and **G**) to old (**F** and **J**). Arrowheads in (**D** to **F**) indicate SBR primordia. Dashed red line in (**H** to **J**) marks the SBR primordia. (**K** to **P**) Confocal images of stem sections of *DR5:mScarleti-NLS TCSn:mNeonGreen-NLS* plants showing auxin (red) and cytokinin (green) responses at the different stages of SBR meristem development from the primary phloem. (**Q**) Single-cell profiling experimental design. Scale bars are 5cm in (**B**), 1cm in (**C** to **F**), 50µm in (**G** to **J**) and 25µm in (**K** to **P**).

### Tomato SBR initiate from differentiated phloem cells in a distinct hormonal dynamic

Under our growth conditions SBR were morphologically observable on the first tomato internode ∼7 days after cessation of internode elongation, but there is plant-to-plant variation in the number of roots (Fig. S1, A and B). As the plants developed, SBR continuously initiate from young internodes, forming a developmental gradient along the stem (Fig. 1, C to J). Our analysis of stem sections revealed that SBR preferentially form at the edge of vascular bundles (Fig. S1, C and D). Anatomical analysis is restricted to later stages of meristem development, when cell proliferation is apparent. To study early events, we generated a transgenic tomato carrying reporters for auxin (DR5 (*25*)) and cytokinin (TCSn (*26*)) transcriptional activity (*DR5:mScarleti-NLS TCSn:mNeonGreen-NLS)*, which allows the simultaneous visualization of both hormonal responses. Before SBR initiation, we observed sporadic expression of *TCSn* in the cambial and phloem region (Fig. 1K). In ∼4-week-old plants, immediately following the cessation of internode elongation, we observed SBR initiation events, marked by induction of both auxin and cytokinin signaling, in a small group of ∼30 cells, clearly located in enlarged primary phloem parenchyma (PP) cells (*27*) (Fig. 1L and Fig. S1A). Subsequently, the expression of auxin and cytokinin separated into distinct domains (Fig. 1, M to O). Interestingly, these hormonal dynamics mirror those occurring during root meristem formation in the embryo (*28*) and differ from those of LR initiation (*3*). When the meristem is composed of ∼5,000 cells and has grown into the surrounding stem cortex, its anatomy and hormonal domains resembled those of a mature root meristem (Fig. 1P and Fig. S1E). We observed similar dynamics in plants carrying the more sensitive auxin response marker *pIAAmotif* (*29*) (Fig. S1, F to J). Based on anatomy and hormonal dynamics, we defined five stages of SBR formation, starting from its initiation from PP cells (Fig. 1, K to P).

### Single-cell profiling of SBR initiation reveals a unique transition state

To understand how PP cells change their identity to form root meristems, we profiled the transcriptome of individual cells from initiating SBR. As SBR-initiating cells are very scarce, they cannot be captured by unbiased profiling methods. Therefore, we used fluorescent hormone response markers to microdissect TCSn*-*marked vasculature prior to SBR initiation, and stages 1, 3 and 5 SBR primordia, followed by cell-wall digestion and isolation of individual fluorescent cells using FACS (Fig. 1Q). To ensure complete characterization of SBR initiation, stage 1 cells were oversampled. Cells were subjected to single-cell mRNA-seq and, following quality control, we obtained 960 cells (230, 399, 89 and 242 cells from the pre-SBR vasculature, stages 1, 3 and 5, respectively).

We used Seurat (*30*) to cluster cells from all stages from the combined dataset. Known marker genes were used to annotate the clusters (Fig. S2, A to F and tables S1 and S2). Consistent with the sorted cell population, we found vasculature lineage identities, such as cambium, procambium and phloem, as well as root cap\initials (Fig. 2A). The distribution of identities varied in the different stages of SBR initiation. Stage 1 SBR were highly enriched with phloem-parenchyma cells, confirming that SBR arise from this tissue. Root cap\initial cells were only found from stage 3 onward, indicating that this distinct identity only develops at later stages of meristem initiation. Surprisingly, we identified a cluster to which we could not assign a clear identity. This cluster, which we named ‘transition’, is highly abundant in stage 1, comprising 37% of the cells in this stage (Fig. 2B). While distinct, transition cells most closely resembled root cap\initials (Fig. S2G). A SlingShot (*31*) trajectory analysis identified two trajectories in the early stages of SBR development, starting from the cambium. One leading to differentiated companion cells and the other, through the transition cells, to root cap\initial cells (Fig. 2C and Fig. S2H), suggesting that the transition cells are progenitors of the new root meristem. The transition identity is characterized by high expression of auxin and cytokinin response genes, as well as expression of mitotic markers (Fig. S3, A to C). Furthermore, this cell type highly expressed ribosomal genes and cell wall invertase, suggesting that it represents a strong carbon sink and has high metabolic activity (Fig. S3, D and E). It was previously suggested that dramatic cell-fate transitions might involve a high-entropy transition state with large transcriptional variability (*32–34*). Indeed, we found that transition cells exhibited high entropy compared to PP or root cap/initial cells (Fig 2D). Overall, our analysis suggests the existence of a previously uncharacterized transient cell identity that is formed in the transition from PP to new root meristem.

**Fig. 2.**
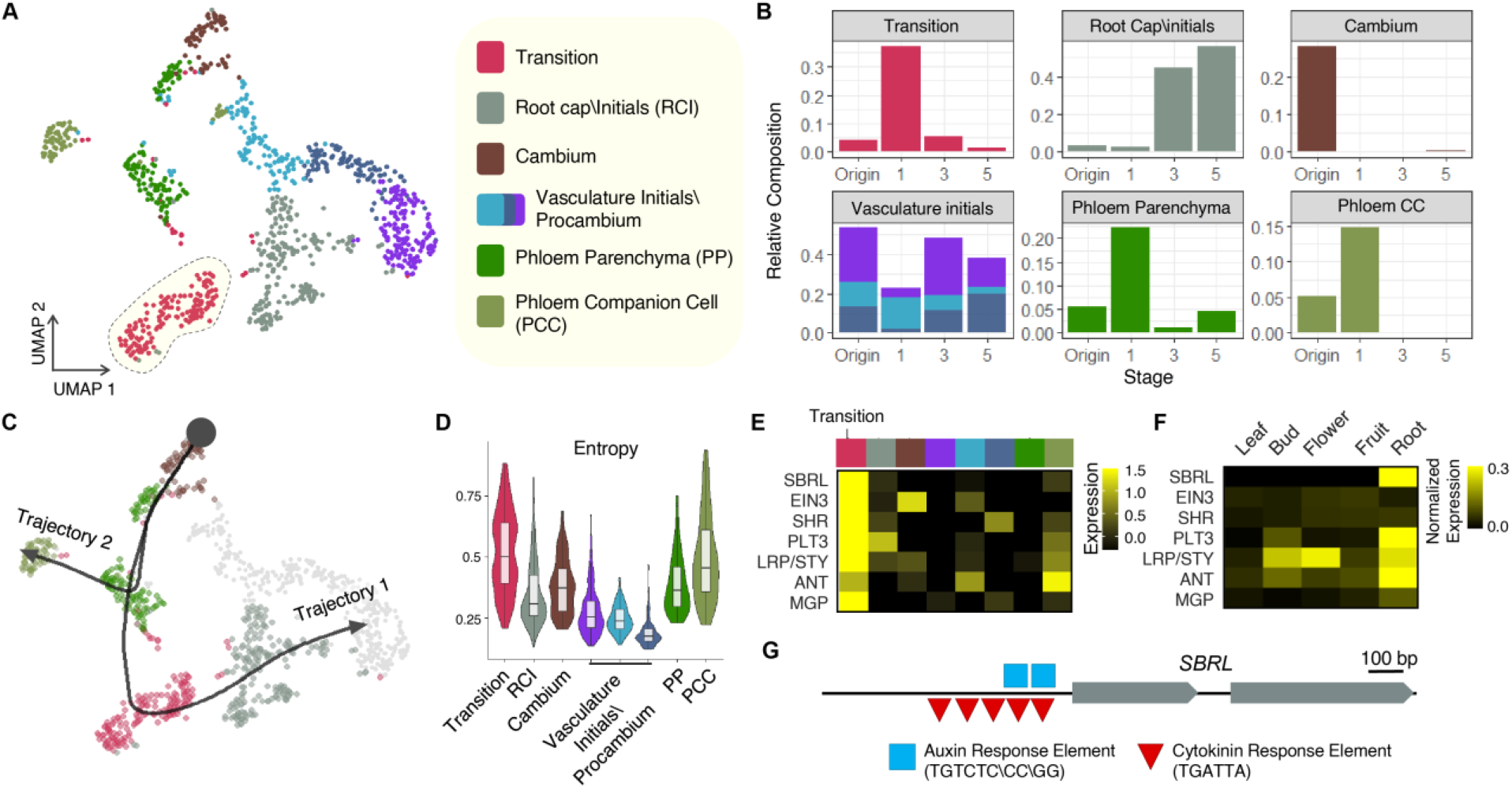
Single-cell transcriptomics of SBR initiation. (**A**) UMAP of cells from SBR development. (**B**) Relative composition of cell identities in each of the four SBR stages. (**C**) Computed trajectory of early stage identities. (**D**) Violin plot of expression entropy for the different identities. (**E** and **F**) Expression of key root development regulators in distinct identities (**E**) and in a global tomato expression profile atlas (**F**). (**G**) Exon (gray bars) structure of SBRL with its auxin and cytokinin response elements, located in the 1.5kb upstream promoter.

### A single transition-identity-expressed LBD gene is required for SBR formation

Among the enriched transcription factors in the transition identity are multiple root stem cell regulators, such as *PLT, SHORT ROOT* and *MAGPIE* (Fig. 2E). Interestingly, *SlWOX11* (*Solyc06g072890*) expression was not detected in our dataset, suggesting that its function may be specific to wound-induced roots (*24*). The transition identity was highly enriched for the expression of a single LBD transcription factor (*Solyc09g066270)*, which we named *SHOOTBORNE ROOTLESS (SBRL*; Fig. S4A). There are 46 LBD genes in tomato, of which *SBRL* is classified as a Class IB LBD (*35*). As members of this class were linked to the regulation of root development (*36*), we decided to further characterize its function. An analysis of a whole-plant expression atlas revealed *SBRL* to have root-specific expression (*37*) (Fig. 2F). *SBRL* promoter has both conserved variants of auxRE and cytRE elements (*29*) (Fig. 2G), and its expression was induced by auxin 6h after treatment in a cytokinin-dependent manner (Fig. S4B).

To verify SBRL expression, we generated *pSBRL:mScarleti-NLS* lines, which confirmed that *SBRL* was induced in stage 1 SBR but not in vasculature tissue (Fig. 3, A and B). We then generated CRISPR mutations of *SBRL* and examined four independent alleles, all of which disrupted the conserved LBD region (Fig. 3C and Fig. S4C). All *sbrl* mutants germinated normally, were fertile and had a normal shoot morphology (Fig. 3, D and E). However, remarkably, all had barren stems that completely lacked SBR, even under flooding conditions that induce SBR emergence in wild type tomato (*38*) (Fig. 3, F to I). To determine whether the defect was in the initiation or emergence of SBR, we generated *sbrl DR5:VENUS-NLS* plants. We were unable to find *DR5-*expressing stage 1 SBR or to identify aberrant cell divisions in phloem tissue in 6 individual plants, suggesting that *SBRL* is specifically required for the earliest stage of SBR initiation (Fig. S4, F and G).

**Fig. 3.**
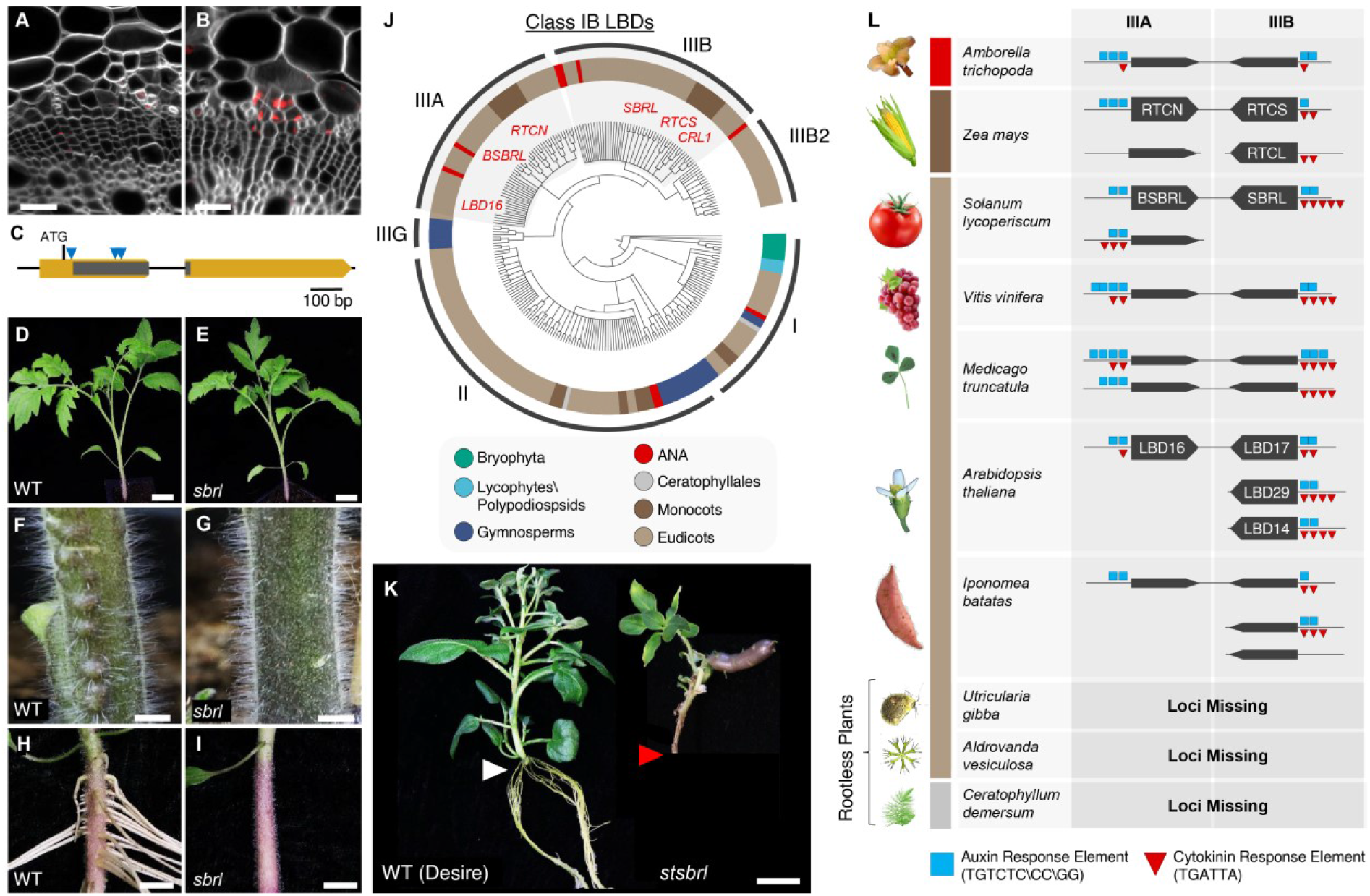
Function and evolutionary conservation of *SBRL*. (**A** and **B**) Confocal images of *pSBRL:mScarleti-NLS-SBRLterm* in vascular tissue (**A**) and at stage 1 SBR (**B**). (**C**) Structure of *SBRL*. Gray bars mark the conserved LOB domain. Blue arrowheads indicate gRNA targets. (**D** to **G**) 4-week-old (**D** and **E**) and a close-up of the first internode of 8-week-old (**F** and **G**) wild type (**D** to **F**) and *sbrl* plants (**E** to **G**). (**H** and **I**), 4-week-old wild type (**H**) and *sbrl* (**I**) tomato hypocotyls after one week of flooding. (**J**) Maximum likelihood tree of class IB LBDs defines six subclasses. (**K**) Wild type and CRISPR-editing *stsbrl* potato plants grown in culture. Note the SBR (white arrow) in the WT, which is missing from *stsbrl* (red arrowhead). *stsbrl* mutants also grew tubers in culture, likely due to systemic effects. (**L**) Synteny and duplication map of subclass IIIA and IIIB genes in selected angiosperms. Scale bars are 25µm in (**A** and **B**), 2cm in (**D** and **E, H** and **I**), and 1cm in (**F** and **G**).

### *SBRL* regulation of SBR is conserved in angiosperms

The complete loss of shoot-borne roots in *sbrl* was reminiscent of the *rtcs* and *crl1* mutations in maize and rice, which are also class IB LBD genes (*20, 21*). To determine whether there is an evolutionary relationship between these genes, we generated a Maximum Likelihood tree from 237 class IB genes obtained from 39 different species (Table S3). This high-resolution analysis revealed that rather than a single class, class IB LBDs form six distinct subclasses. Subclass I was found in all plants, including basal plants, and likely represents the original member of the class. Subclass II appeared in gymnosperms and angiosperms. Subclass IIIG was unique to gymnosperms, while subclasses IIIA and IIIB/IIIB2 were only found in angiosperms. *SBRL, RTCS*, and *CRL1* all belong to subclass IIIB (Fig. 3J).

The vasculature anatomy and SBR physiology differ greatly between dicots and the distant monocots (*39*). Thus, it was surprising that a mutation in a single subclass IIIB gene led to the loss of SBR in all three species. This raised the hypothesis that the regulation of SBR initiation by subclass IIIB genes may be deeply conserved in plants. To test this, we examined the expression of *SBRL* orthologs in stems of white beans (*Phaseolus vulgaris*), sweet potato (*Ipomoea batatas)* and cucumber (*Cucumis sativus*), all of which can naturally produce *SBR* (Fig. S5, A to C). A qPCR analysis revealed that the expression of subclass IIIB genes was induced in stems of all three species immediately prior to the appearance of SBR (Fig. S5E).

To determine whether the function of IIIB orthologs is conserved, we generated CRISPR mutants in the potato (*Solanum tuberosum*) ortholog of *SBRL, StSBRL* (PGSC0003DMG400009716; Fig. S6, A and B). In contrast to wild type controls, three independently derived CRISPR mutants did not produce SBR when propagated in culture and as a result failed to root and could not be maintained (Fig. 3K). A promoter analysis for subclass IIIB genes from different species revealed similar promoter structure with conserved auxin and cytokinin response elements (Fig. 3L and Figs. S5D and S6A), consistent with previous reports that found highly significant hormonal response element conservation for the subclass IIIB gene *LBD17* in *arabidopsis* (*29*). Taken together, this suggests that the SBR initiation program and its regulation by subclass IIIB LBDs is deeply conserved in angiosperms.

### A conserved superlocus regulates root initiation

During the assembly of the phylogenetic tree, we observed that, despite the rapid expansion of the class IB genes (Table S3), subclass IIIB genes were always located immediately next to a closely related subclass IIIA gene in a single inverted duplication superlocus. We identified just three species that lacked this superlocus, two carnivorous plants (*Aldrovanda vesiculosa* and *Utricularia gibba*) and the water plant *Ceratophyllum demersum*). Common to all three is that they lost their roots during evolution (Fig. 3L). The conserved physical proximity and sequence homology of subclass IIIA and IIIB genes suggests that they may play redundant function in root initiation. However, the tomato subclass IIIA *BROTHER OF SBRL* (*BSBRL*), located immediately adjacent to *SBRL*, was not expressed at all in the SBR single-cell data. Instead, two other subclass IIIA genes, the maize *RTCN* and the *Arabidopsis LBD16*, were previously linked to the regulation of LR initiation (*11, 13, 14, 40*), suggesting that subclass IIIA genes may play a conserved role in LR initiation. To test this hypothesis, we first characterized LR initiation in tomato.

Similar to *Arabidopsis* LR initiation, and in direct contrast to SBR initiation, LR starts with two pericycle cells activating the auxin response, flanked by cells expressing the cytokinin response marker (Fig. 4, A and B). Auxin response then marks the forming primordia until it reaches its mature form (Fig. 4, C to F). To test whether a transition state similar to that observed during SBR formation occurs during the initiation of lateral roots, we divided LR initiation to six stages and generated RNA-seq profiles of microdissected tomato LR from each one (Fig. 4, A to F). Using the average expression of identity markers from the single-cell analysis (Table S2), revealed that markers of the transition state were induced at stage 1, followed by markers of root-cap/initial identities and then procambium markers, suggesting a similar transition identity may occur briefly during LR initiation (Fig. 4G). Both genes in the superlocus (*SBRL-BSBRL)*, as well as another duplicate of subclass IIIA (Fig. 3L), were upregulated at the earliest, two-cell stage of LR initiation, expressed throughout meristem formation, but downregulated at later stages (Fig. 4H). To test whether the co-activation of subclass IIIA and IIIB in LR initiation is a conserved feature in plants we measured the expression of superlocus genes in isolated root maturation zone of *P. vulgaris* and *C. sativus*. Similarly to tomato, both subclasses were induced during LR initiation, but only subclass IIIB were induced during SBR initiation in both species (Fig. 4I and Fig. S5E). This is consistent with previous reports in maize showing that subclass IIIA gene *RTCN* is highly enriched in LR, while the expression of subclass IIIB gene *RTCS* is confined to shoot-borne nodal roots (*20*). Our analysis of the *pSBRL:mScarleti-NLS* reporter confirmed the transcriptome dynamics and revealed that, similarly to the *Arabidopsis* subclass IIIB gene *LBD29* (*41*), *SBRL* is also expressed in the endodermis and cortical cells overlaying the emerging LR (Fig. 4J to M).

**Fig. 4.**
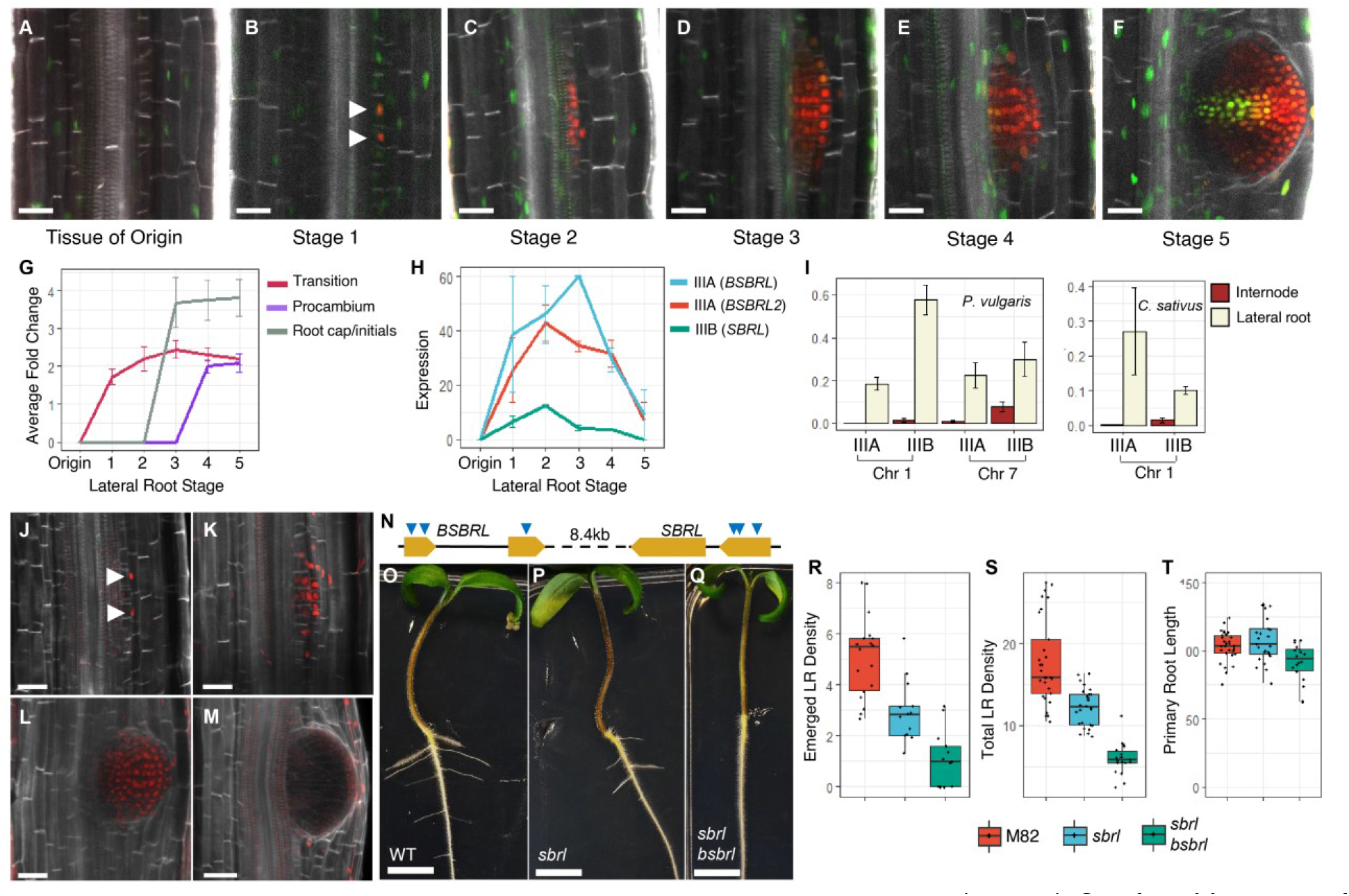
A conserved superlocus regulates lateral root initiation. (**A** to **F**) Confocal images of *DR5:mScarleti-NLS TCSn:mNeonGreen-NLS* before (**A**) and during (**B** to **F**) LR development. (**G**) Average induction of identity markers during LR development. (**H**) The expression of subclass IIIA and IIIB LBDs during LR development. (**I**) qRT-PCR of subclass IIIA and IIIB LBDs in *P. vulgaris* and *C. sativus* in lateral roots compared to young internodes (error bars are the standard error of 4 biological replicates). (**J** to **M**) Confocal images of a temporal series of *pSBRL:mScarleti-NLS-SBRLterm* roots during LR initiation. (**N**) Structure of the *SBRL-BSBRL* superlocus. Blue arrowheads mark the targets of the multiplex CRISPR gRNA. (**O** to **Q**) Representative picture of 10-day-old wild type (**O**), *sbrl* (**P**) and *sbrl bsbrl* (**Q**) tomato seedlings. (**R** to **T**) Emerged LR density (**R**), combined emerged and pre-emerged LR density (LR/10cm) (**S**) and primary root length (mm) (**T**) in 10-day-old wild type, *sbrl* and *sbrl bsbrl* mutants (n=29,26 and 18, respectively). The total LR density of both *sbrl* and *sbrl bsbrl* is significantly reduced compare to WT (pval=1E-6 and <1E-30, respectively; Tukey HSD). Arrowheads in (**B**) and (**J**) points to the two LR initial cells. Scale bars are 50µm in (**A** to **F** and **J** to **M**) and 1cm in (**O** to **Q**).

As both subclass IIIA and IIIB are coexpressed during the first stage of LR initiation, we hypothesized that they serve a redundant role in the process. We used multiplex CRIPSR with 6 guides to target the *SBRL-BSBRL* superlocus (Fig. 4N) and examined four independently derived alleles (Fig. S7A), all showing the same phenotype. Remarkably, *sbrl bsbrl* double mutants exhibited a dramatic reduction in emerging and pre-emerging LR primordia, while the primary root and shoot remained unaffected (Fig. 4, A to T).

Overall, we show that an ancient duplication in a class IB LBD gene, which occurred early in angiosperm evolution, resulted in rapid subfunctionalization of the duplicate genes, with one gene assuming specific control of SBR initiation. We suggest that this locus and the genes subsequently spawned from it allow the separation of environmental inputs guiding the initiation of different root types and, thus, the remarkable diversity in root/shoot relationships found among vascular plants (*5*).

It has been hypothesized that roots that form in different contexts differ in their early ontogeny, but converge on a similar genetic program (*42*). Indeed, we show that LR and SBR differ in terms of the number of initial cells involved and their hormonal dynamics, while their morphologies become similar at later stages. Early LR are made of just two cells and their limited accessibility makes single-cell level profiling of this stage extremely challenging. However, single-cell expression data from cells forming a root stem-cell-niche following injury revealed a transient state that is similar to the transition identity (*43*), suggesting that an early high-activity transition identity may be a general feature of *de novo* meristem formation.

## Acknowledgments

We thank Tom Beeckman, Yuval Eshed and Sigal Savaldi-Goldstein for comments and discussions. Shmulik Wolf for materials and Itamar Pri-Tal for assistance in tissue culture.

## Author contributions

M.O., N.G-Y. and I.E. conceived and designed the study and performed the experiments; C.Y. performed the SBR quantification and LR RNA-seq assay; E.S. performed the potato transformation; I.E., N. G-Y. and M.O. wrote the manuscript.

## Data and materials availability

Raw and processed data is available in GEO: Single cell (GSE159055), lateral root transcriptome (GSE159050).

## Supplementary material and methods

### Plant materials and growth condition

Tomato *(Solanum lycopersicum* cv M82) seeds were germinated on potting soil, grown in a growth chamber in a 16/8 light/dark regimen at 24°C for 6 weeks and then transplanted to 10L pots to continue their growth in a climate controlled greenhouse (natural day length, 25°C /20°C day/night temperature) for additional 4-6 weeks. *DR5:3xVENUS-NLS* and *TCS:3xVENUS-NLS* plants were described before(*44, 45*). Common bean (*Phaseolus vulgaris*) and Cucumber (*Cucumis sativus* cv. Kfir) were germinated grown in a closed moist chamber at room temp. At 10 days after germination plants were transfers to 5L pots and grown in the greenhouse under the same conditions. Sweet potato (*Ipomoea batatas cv ‘Georgia Jet’*) for RT-PCR were grown from tuberous roots in water at room temperature under ambient light.

### Transgenic plants

All cloning steps were performed using the MoClo system(*46*) with synthesized DNA fragments. For the construction of the dual hormone reporter line we first generated level 0 constructs. The DR5 and TCSn sequences(*26, 47*) were cloned into *pICH41295. mScarleti* and *mNeonGreen* were codon optimized for plants and cloned into the *pAGM1287*. The NLS sequence N7 was cloned to pAGM1301. rbcS-E9t and HSP18.2 terminators were synthesized and cloned into *pICH41276*. To assemble the construct, DR5, mScarleti, N7 and rbcS-E9t were cloned into level 1 vector *pICH47742*, and TCSn, mNeonGreen, N7 and HSP18.2 terminator were cloned to level 1 vector *pICH47751*. Both assembled level 1 clones were assembled to the binary vector *pICSL4723* carrying a Kanamycin resistance cassette. To produce the high sensitivity auxin marker, the synthetic auxin reporter pIAAmotif(*29*) was cloned to level 0 vector *pICH41295*. Level 1 was assembled by cloning the pIAAmotif, mScarleti, N7 and rbcS-E9t terminator into *pICH47742*, which was then assembled to a binary level 2 vector *pICSL4723*. For production of the *SBRL* reporter, its 2.6 Kb upstream sequence was amplified in two fragments and assembled to level 0 vector *pICH41295*, and its 0.6Kb downstream sequence (termination sequence) was amplified and assembled to level 0 vector *pICH47761*, which were together used to build a level 1 construct with mScarleti, and N7 in *pICH47751*. Level 1 was then assembled to a level 2 binary *pICSL4723* with Kanamycin resistance.

For CRISPR/Cas9 constructs, gRNA were prepared as previously described(*48*). The 2kb sequence of tomato ubiquitin 10 promoter (*SlUBQ10/Solyc07g064130*) including the first intron was cloned into the level 0 vector *pICH41295. Zea mays* codon-optimized Cas9 (zCAS9) was cloned to pAGM1287. Level 1 was assembled by cloning the tomato ubiquitin 10 promoter, zCAS9, and rbcS-E9t terminator into *pICH47742*. CAS9 Level 1 vector was then assembled to the level 2 binary *pICSL4723* with Kanamycin resistance. gRNA target sites including the PAM site (NGG) were designed against the tomato genome (Solanum lycopersicum str.Heintz 1706) using Benchling (https://www.benchling.com). gRNAs target sites were selected manually, based on their on-target score^4^. gRNAs were subsequently assembled into the binary Level 2 vector carrying CAS9 and Kanamaycin resistance.

### Plant transformation

Level 2 binary vectors were introduced into the *Agrobacterium tumefaciens* strain GV3101 by electroporation. For tomato, transgenic plants were generated by cotyledon transformation as described(*49*). Potato (*Solanum tuberosum* cv. Desirée) leaf explant were transformed as described(*50*). Once the regenerated shoots reached 2-3 cm long, they were cut at the base and transferred to jars containing MS medium (Duchefa; MO222.005) supplemented with 0.8% agar, 1.6% glucose, 250mg/L claforan, 50mg/L kanamycin, and were grown under long day (16hr/8hr) conditions at 22°C. Three independent homozygous lines were cut and transferred into new jars and grown together with WT plants. The WT plants produced roots after two weeks, while the *sbrl* mutant lines did make roots even after 8 weeks.

### Characterization of SBR and LR development in tomato

4-6 weeks-old M82 plants were harvested to analyze the developmental stages of SBR formation and observed by stereomicroscope. SBRs were scored by the appearance of a discoloration on the stem than precedes root emergence. For LR analysis, tomato seeds were sterilized by incubating in 75% (v/v) ethanol for 3 min, followed by bleach solution 3% (v/v) with 100 μl/l Tween 20 for 15-20 min. Seeds were washed in sterile water and germinated on 12.5cmx12.5cm agar plates containing 1/2 Murashige and Skoog medium (Sigma M5519), 0.5% sucrose, 0.8% agar at pH 5.7. At 10 days after seeding, LR staging was performed under a stereoscope. Emerged LR were those that grew beyond the epidermis. Pre-emerged LR were identified by the localized growth from the pericycle. Root length was measured using ImageJ.

### Confocal Imaging

Fresh stems transverse sections of internode 1 were hand-sectioned with a razorblade, observed under the Nikon SMZ18 fluorescent stereomicroscope for the presence and SBR staging, followed by fixation and clearing using ClearSee^6^. Cleared sections were stained with SCRI Renaissance 2200 (SR2200, 0.1% (v/v)) for in ClearSee solution for 2 to 72 hours, mounted in ClearSee solution and scanned with Leica SP8 confocal microscope. Solid state lasers were using for excitation (488nm for Venus and mNeonGreen; 552nm for mScarleti and 405nm for SR2200).

### Cell separation and FACS

For cell isolation, internode 1 was slices to thin slices using a razor blade and SBR primordia were isolated based on DR5 or TCS expression under a fluorescence stereoscope. Cortex and epidermis tissues were removed and SBR primodia microdissected using fine 18G needles. Ttissue sections were transfers to cell wall digestion solution (3% w/vol Cellulase, 1% w/vol Macerozyme, 0.4M Mannitol, 20.48mM Mes, 0.02M Kcl, Tris-Hcl pH=4.5, Heated to 55C for 10min, cooled to room temperature and add 0.001gr/ml BSA, 20mM CaCl2). Following 15 min of gentle mixing in the cell wall digestion solution, slices were pipetted carefully with a cut tip and returned for an additional 15-45 min of cell wall digestion. Solution was then filtered using 40 micron filter, stained using SR2200 for cell debris. Cell separation was done using FACS Aria or FACS Melody. Cells were gated according to negative SR2200 on V450 channel and positive for neon green on FITC channel. Cell separation was done to a 96 wells plate containing 5µl lysis buffer.

### Single-cell mRNA-Seq

Amplification of single cell RNA was performed according to the mc-SCRB protocol(*51*). cDNA libraries quality was tested using PCR for the presence of *ELONGATION FACTOR 1* (*EF1*) and *VENUS/mNeonGreen* transcripts. Sequencing libraries were constructed using Nextera XT (FC-131-1024) from 1.2ng of amplified cDNA. During library PCR, 3’ ends were enriched with a custom P5 primer (P5NEXTPT5, IDT). Libraries were pooled, size-selected using 2% Agarose Gels to remove primer concatamers and gel-extracted using the MinElute Kit (Qiagen) according to manufacturer’s recommendations. Paired-end sequencing (16bp for cellular barcode and UMI; 50bp for cDNA). Following sequencing, FASTQ files were filtered for poly-A reads using a custom script and aligned to the ITAG4 genome using STAR 2.7.1. To account for the low accuracy of 3’UTR annotation in tomato, all genes were extended by 500bp. Read calling was performed using the zUMI pipeline(*52*). Analysis were done using the Seurat package v3(*30*) with R v4.0.1. Genes that appear in less than 4 cells or have less 20 UMIs in the entire dataset were removed. Then, cells having less than 400 genes or 10,000 UMIs, or more than 7000 genes or 30,000 UMIs were filtered. Data was collected from 11 independent biological batch, each containing a mixture of SBR stages. Two methods were used to correct for batch effects. First, we calculated the contribution of batch to gene variance and 415 genes with values over 0.15 were removed. Then, the CCA pipeline (RunCCA) for batch correction with 2000 variable genes was used to further correct for batch effect. To visualize the data, we performed principal component analysis (PCA) for dimensionality reduction using 800 variable features, followed by UMAP with the first 15 PCA dimensions. Clusters were identified using the same parameters using the seurat FindClusters function (parameters: resolution = 0.7, algorithm = 2, n.start = 1000, n.iter = 10000).

Pseudotime trajectory analysis was performed using the SlingShot R package algorithm (40) using top 100 variable genes. The analysis was performed with the UMAP dimensionality reduction and cluster labels as in Seurat objects to identify the trajectory. For clarity and brevity, multiple trajectories were plotted on the same graph. Calculation of cell gene expression entropy were performed by binning expression into 100 discrete bins, followed by application of the *entropy* function of the *entropy* R library. The auxin, cytokinin and ribosomal response indices were calculated as the average expression of the auxin-responsive *SlIAA1/2/3/4* (Solyc09g083280, Solyc06g053840, Solyc09g065850, Solyc06g084070), the cytokinin-responsive A-class ARR (Solyc02g071220, Solyc03g113720, Solyc05g006420, Solyc06g048930, Solyc10g079600, Solyc10g079700) or of annotated ribosomal genes, respectively.

### Phylogenetic analysis

Genome and proteome sequences were obtained from phytozome (https://phytozome.jgi.doe.gov/), with the exception of sweet potato (*Ipomoea batatas)* genome v3, obtained from the Ipomoea Genome Hub (http://www.ipomoea-genome.org/); *Arabis alpina* genome(*53*) v4, obtained from (http://www.arabis-alpina.org/); *Aldrovanda vesiculosa* genome(*54*),obtained from (https://www.biozentrum.uni-wuerzburg.de/carnivorom/resources) and *Vigna radiata* v1 from the Legume Information System (https://legumeinfo.org/). Class IB sequences were by searching the each proteome for the canonical LBD class IB sequence CGACKFLRRKC(*36*). To identify unannotated LDB genes, BLASTX was used to search the canonical class IB LDB sequence in the genome. The LBD domain sequences of the proteins (108-127aa in length) was aligned using MUSCLE. Phylogenetic tree was constructed using the RAxML package version 8(*55*) using the GAMMAJTT model and with 1,000 bootstrap runs (raxmlHPC-PTHREADS-SSE3 -T 16 -f a -x 12345 -N 1000 -p 12345 -m PROTGAMMAJTT). Tree visualization was performed using iTOL. Branches with bootstrap value of less than 30 were collapsed.

### Lateral Root developmental series

*DR5:3xVENUS-NLS* tomato plants were germinated on agar as described above. At 7 days root were observed under a fluorescent stereoscope to identify LR stage. Thin root sections contain the LR at appropriate stage were dissected and flash frozen. 4-6 tissue sections were used for each sample. Two independent biological samples were collected for each stage. RNA extracted using Qiagen RNeasy Micro Kit (cat. 74004), libraries were concentrated using QuantSeq 3’ mRNA-Seq Library Prep Kit (LX-015.96) and sequenced using NextSeq 500 with 75bp single read reads. Expression calling was performed using Salmon v1.3.0(*56*) with tomato ITAG4 transcriptome. All transcripts were extended by 500bp. Expression calls with less than 3 reads were discarded. Normalization and differential expression was performed using DESeq2 v1.28.1(*57*)

### qRT-PCR

For measurement of class IB LBD expression, RNA was isolated from the following tissues: *I. batatas:* 0.5cm section of the stem-leaf junction were collected from 3 successive nodes with the last node have clear SBR growth. Junctions were collected from three individual plants. *C, sativus* and *P. vulgaris:* 2cm sections of 3 consecutive internodes were collected from 3 individual plants. Not all plants of this species produced SBR and there was plant-to-plant variability. For lateral roots analysis, a 1cm section from ∼1cm about the meristem was collected in groups of 4 roots per sample. Four independent biological replicates were collected for each species. For measurement of hormonal response in tomato stems, hormone solutions were gently injected into internodes 1 of 4 weeks-old tomato plants using 2.5 ml syringe with 30G hypodermic (yellow) needle. Internode sections were collected 6h after injection. All tissues were flash frozen and RNA extracted using Trizol (Bio-Lab 009010233100). Total RNA was treated using TURBO DNA-free™ Kit (Thermo Fisher) and cDNA prepared using qScript CDNA kit (QuantaBio), according to the manufacturer’s instructions. Real-time PCR was performed on qTower 3 thermal cycler using Fast SYBR Green Master Mix (Rhenium) with a final primer concentration of 0.2μM. Reference genes were UBQ for *S. lycoperiscum*, IDE for *P. vulgaris*(*58*), Actin for *I. batatas*(*59*), and *CACS* for *C. sativus*(*60*). Primers are listed in (table S4).

**Fig. S1.**
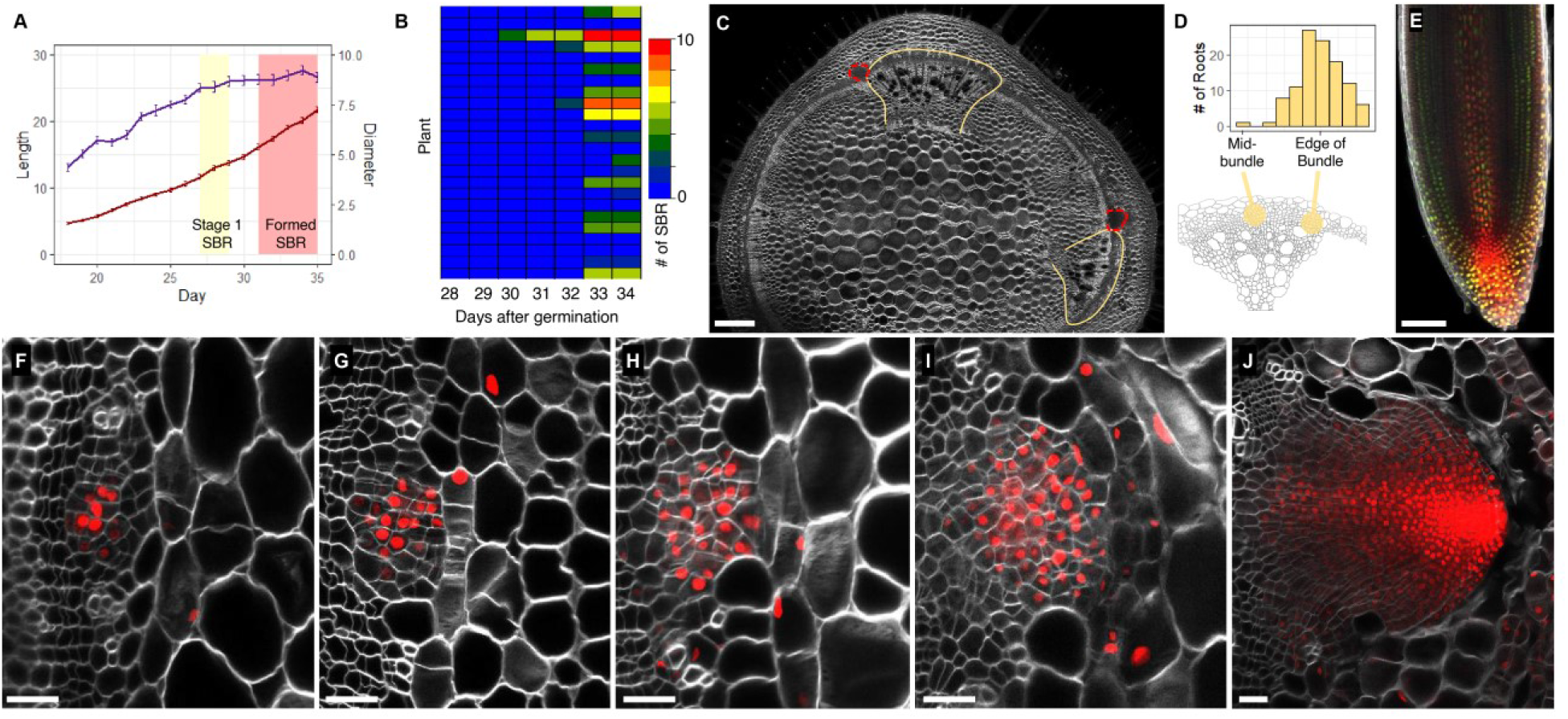
Dynamics of tomato SBR formation. (**A**) Length (purple) and width (red) of internode 1. (n=24). The shading marks the approximate day when stage 1 (yellow) and fully formed (red) SBR primordia appear. (**B**) Timing and number of SBR on internode 1 of 24 individual tomato plants grown under similar conditions. (**C**) Confocal image of an entire stem cross-section of a 34-day-old plant. Yellow lines mark the vascular bundles. Red dashed lines indicate forming primordia. (**D**) The distribution of the SBR formation position relative to the vascular bundle. (**E**) Confocal picture of *DR5:mScarleti-NLS TCSn:mNeonGreen-NLS* of a primary root tip. (**F** to **J**), Confocal images of a temporal series of SBR initiation in *pIAA:mScarleti-NLS* plants (from left to right). Scale bars are 100µm in (**C** to **E**) and 25µm in (**F** to **J**).

**Fig. S2.**
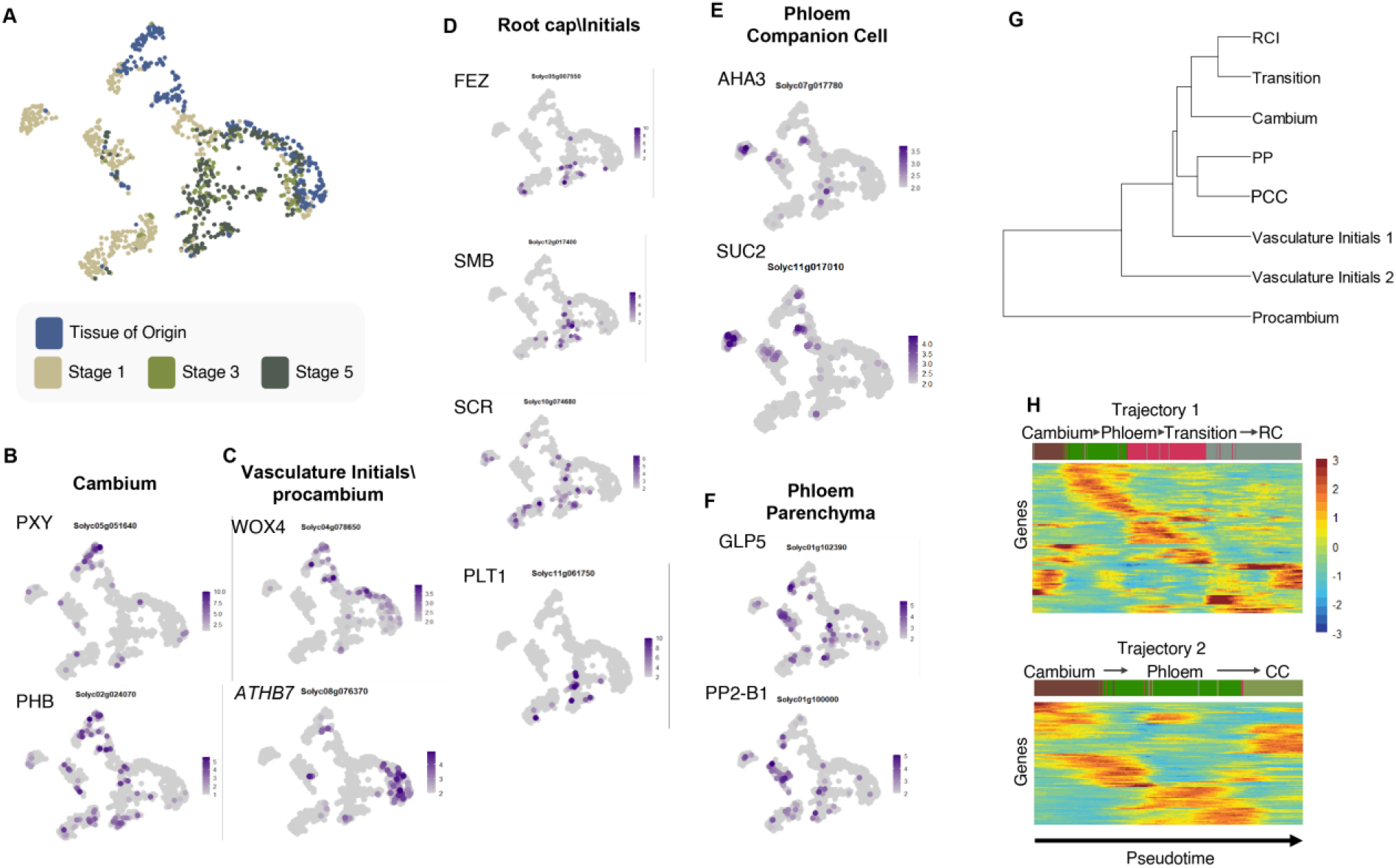
Stage- and tissue-specific expression during SBR development. (**A**) UMAP of single cells coded by developmental stage. (**B** to **F**) Expression of known identity markers. (**G**) Cluster tree showing the similarity between different cell identities. RCI, root cap\initials; PP, phloem parenchyma; PCC, phloem companion cells. (**H**) Heatmap of gene expression along the two early trajectories predicted by Slingshot.

**Fig. S3.**
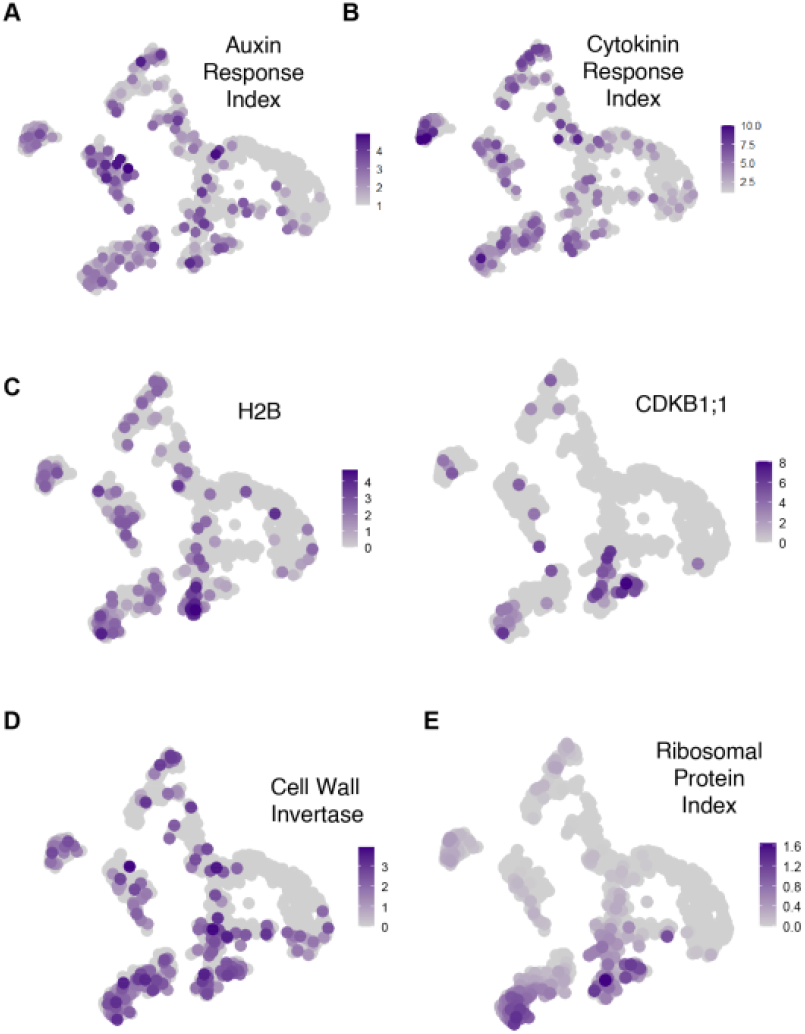
Expression patterns in the transition identity. (**A** to **E**) The expression of auxin (**A**) and cytokinin (**B**) response indices, cell cycle markers (**C**), cell wall invertase (**D**) and ribosomal protein index (**E**) in single cells.

**Fig. S4.**
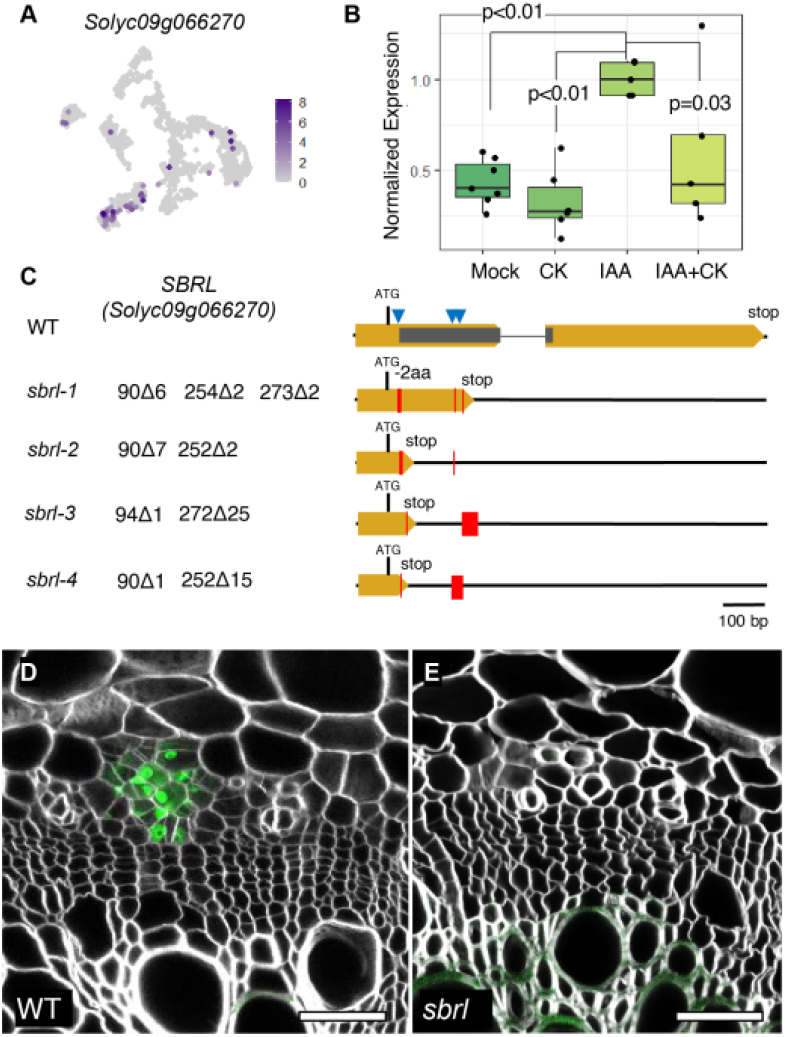
*SBRL’s* role in root initiation. (**A**) The expression of *SBRL* in the single cell data. (**B**) qPCR expression of *SBRL* in internode 1 of 4 weeks old tomato 6h after being injected with mock, 25µM kinetin (CK), 5µM IAA or both IAA and CK, normalized to expression of IAA treated plants (n=7,6,5,5 for mock, CK, IAA and CK+IAA treated, respectively. p-values are Tukey HSD for GLM). *SBRL* expression was induced by IAA, but this induction was suppressed by co-treatment with CK. (**C**) The *SBRL a*lleles used in this study. Blue arrowheads indicate gRNA target sites. Red bars indicate deletions. (**F** and **G**) Stem sections of *DR5:3xVENUS-NLS* (**F**) and *sbrl DR5:3xVENUS-NLS* (**G**). No *DR5* peaks were observed in any of the *sbrl* stems (n=6). Scale bars are 25µm.

**Fig. S5.**
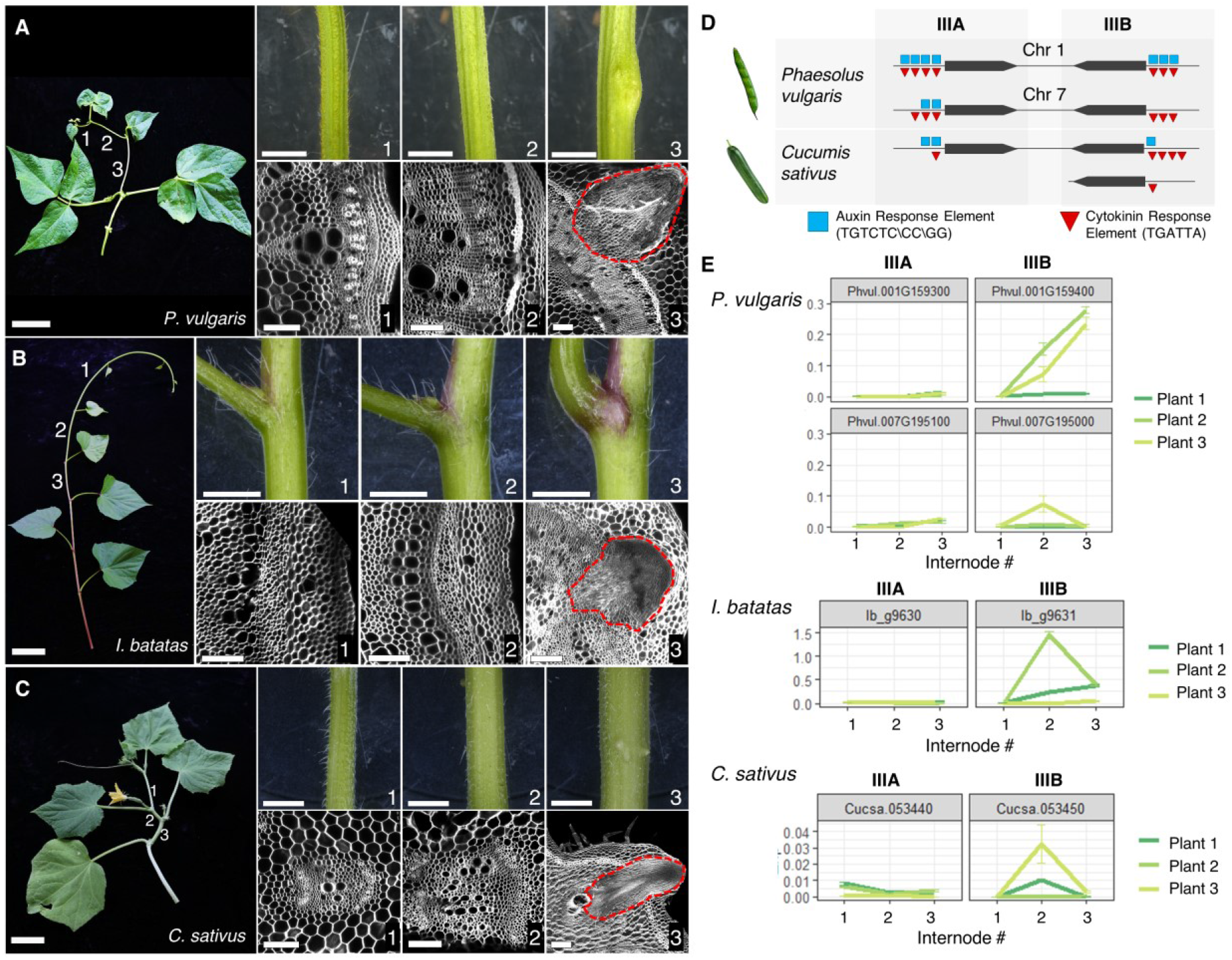
SBR development and Class IB LBD expression in different species. (**A** to **C**) Whole plant (left, scale bars are 5cm), closeup of specific internodes (top, scale bars are 0.5cm) and matching confocal images of the stem section (bottom, scale bars are 100µm) for *P. vulagris* (**A**), *I. batatas* (**B**) and *C. sativus* (**C**). Red dashed line marks the SBR. (**D**) Genetic structure of subclass IIIA\IIIB genes in *P. vulgaris* and *C. sativus*. (**E**) The expression of subclass IIIA and IIIB genes in consecutive internodes of 3 individual plants of *P. vulgaris, I. batatas* and *C. sativus*. As the rate of SBR formation varies from plant to plant, each plant was assayed individually. Error bars represent the standard error of 3 technical replicates.

**Fig. S6.**
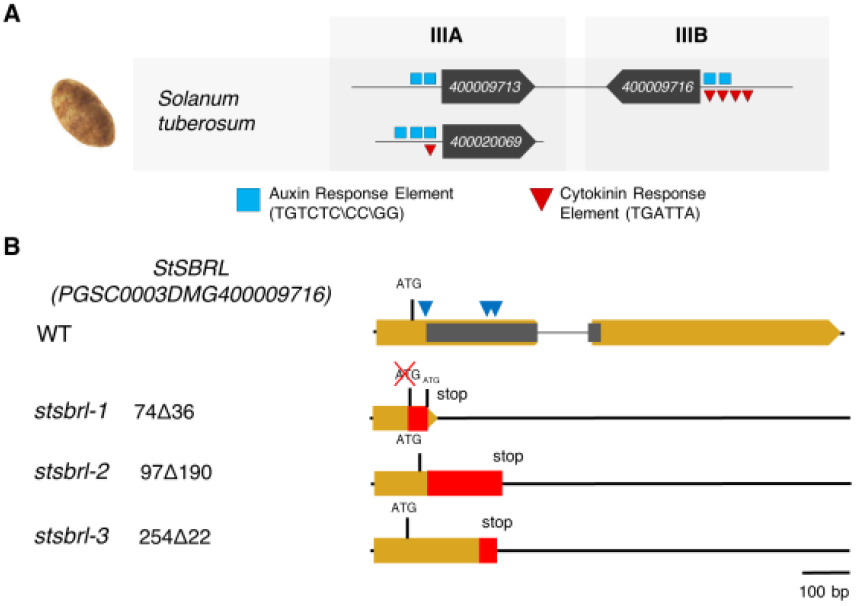
Class IB LBD genes in potato. (**A**) Genetic structure of subclass IIIA\IIIB genes in *S. tuberosum*. Gene names are noted on the locus. (**B**) The genetic lesions in potato *StSBRL* alleles used in this study. Blue arrowheads indicate gRNA target sites. Red bars indicate deletions.

**Fig. S7.**
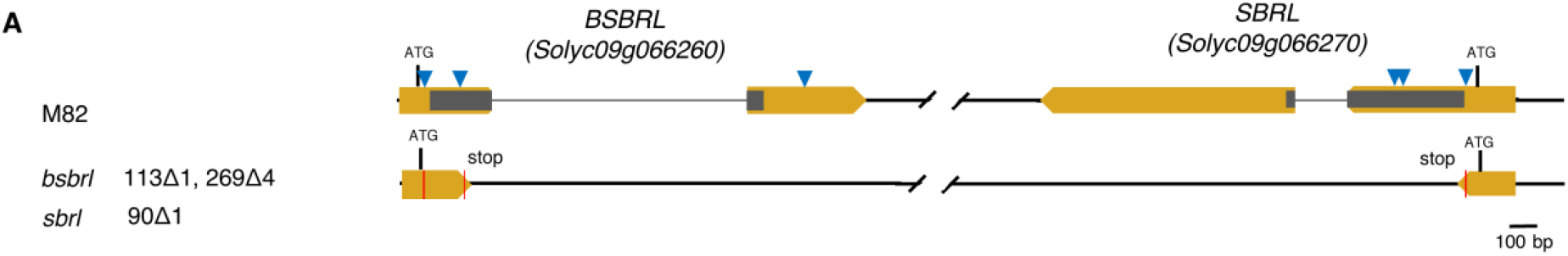
CRISPR-generated mutant alleles of the *SBRL-BSBRL* superlocus. (**A**) Genetic lesions in four independently derived T0 tomato plants. All four independent CRISPR lines had identical mutations.

**Supplemental Table 1. List of known tissue-specific marker genes used to annotate single-cell clusters**

**Supplemental Table 2. List of identity-specific marker genes**

**Supplemental Table 3. Class IB LBD genes**

**Supplemental Table 4. Primers used in this study**

